# Expedited gene delivery for osteochondral defect repair in a rabbit knee model: a one-year investigation

**DOI:** 10.1101/2021.09.22.461300

**Authors:** Christopher V. Nagelli, Rodolfo E. De La Vega, Michael Coenen, Consuelo Padilla de Lopez, Joseph A. Panos, Alejandro Tovar, Sebastian A. Müller, Christopher H. Evans

## Abstract

**Objective:** To evaluate a single-step, gene-based procedure for repairing osteochondral lesions.

**Design:** Osteochondral lesions were created in the patellar groove of skeletally mature rabbits. Autologous bone marrow aspirates were mixed with adenovirus vectors carrying cDNA encoding green fluorescent protein (Ad.GFP) or transforming growth factor-β_1_ (Ad.TGF-β_1_) and allowed to clot. The clotted marrow was press-fit into the defects. Animals receiving Ad.GFP were euthanized at 2 weeks and intra-articular expression of GFP examined by fluorescence microscopy. Animals receiving Ad.TGF-β_1_ were euthanized at 3 months and 12 months; repair was compared to empty defects using histology and immunohistochemistry. Complementary in vitro experiments assessed transgene expression and chondrogenesis in marrow clots and fibrin gels. In a subsequent pilot study, repair at 3 months using a fibrin gel to encapsulate Ad.TGF-β_1_ was evaluated.

**Results:** At 2 weeks, GFP expression was seen at variable levels within the cartilaginous lesion. At 3 months, there was a statistically significant improvement in healing of lesions receiving Ad.TGF-β_1_, although variability was high. At 12 months, there was no difference between the empty defects and those receiving Ad.TGF-β_1_ in overall score and cartilage score, but the bone healing score remained higher. Variability was again high. In vitro experiments suggested that variability reflected variable transduction efficiency and chondrogenic activity of the marrow clots; using fibrin gels instead of marrow provided more uniformity in healing.

**Conclusions:** This approach to improving the repair of osteochondral lesions holds promise but needs further refinement to reduce variability and provide a more robust outcome.

## INTRODUCTION

Trauma to articular cartilage is found in 60% of knee arthroscopies.^1^ A majority of lesions include injuries to both the cartilage and subchondral bone (osteochondral defects).^1^ These defects do not heal spontaneously and, if left untreated, can cause pain and dysfunction leading to post-traumatic osteoarthritis. Cartilage has no innate ability to repair, because the tissue is avascular, aneural, alymphatic, and does not contain stem or progenitor cells.^2^ This has led to cartilage repair surgery being the current clinical standard of care for patients with chondral lesions; surgeons perform approximately 500,000 cartilage repair procedures annually in the US.^3^

Autologous chondrocyte implantation (ACI) and microfracture are two popular cell-based therapies used by surgeons to treat chondral lesions. ACI has shown great promise as a cartilage repair procedure. However, this is a two-step procedure which involves harvesting cartilage from the periphery of the joint, expanding the isolated chondrocytes in culture, and transplanting these cells back into the defect to regenerate tissue.^4^ This increases the cost of the procedure, and patients incur a prolonged post-operative rehabilitation period during which full weight bearing is delayed.^5 6^ In contrast, microfracture is a one-step, point of care procedure which involves exposing and penetrating the subchondral plate underneath the cartilage defect to allow bone marrow to enter and form a fibrin clot within the lesion.^7^ The coagulated bone marrow contains mesenchymal stromal cells (MSCs) with the potential to differentiate into chondrocytes which initiate the repair process.^7^ Microfracture has demonstrated promising short-to-medium term relief for many patients allowing them back to previous levels of activity.^8^ However, according to multiple long-term reports both ACI and microfracture have high failure rates and patients are likely to develop repair tissue resembling a fibrocartilaginous scar instead of native articular cartilage.^9–11 12–14^ Fibrocartilage has inferior biomechanical properties and does not endure the joint environment. Its deterioration causes pain, dysfunction, and eventual failure of the repair, requiring salvage surgery.

The present study is based on the hypothesis that the long-term clinical outcome of microfracture and related marrow stimulation techniques would be improved if the MSCs differentiated fully into articular chondrocytes instead of fibrochondrocytes. Morphogens, such as TGF-β_1_, hold promise in this respect but they are difficult to deliver to osteochondral lesions in a sustained fashion. Gene transfer offers a technology for overcoming this barrier, and we have shown that MSCs transduced with adenovirus vectors encoding TGF-β_1_ undergo efficient chondrogenesis.^15, 16^ Based upon these considerations, the research described in this paper investigates a gene therapy approach for osteochondral defect repair.

We have developed a technology in which autologous bone marrow coagulates incorporating adenoviral vectors are used for gene transfer to osteochondral defects.^17^ The MSCs within the coagulate are transduced by the adenovirus, and the fibrin scaffold retains additional vector for transducing MSCs as they enter the lesion. The marrow clot also has excellent handling properties and conforms to the dimensions of the structure within which it clots. Sieker et al^18^ have published promising results using the marrow clot technology in conjunction with adenovirus encoding bone morphogenetic protein 2 (BMP-2) and Indian hedgehog (IHH) transgenes. While successful in the short term, defects receiving BMP-2 progressively formed bone, while the cartilage in those receiving Indian hedgehog was immature and the subchondral bone absent.

In the present study, we investigated whether delivering adenoviral vectors carrying TGF-β_1_ cDNA (Ad.TGF-β_1_) using autologous bone marrow coagulates can improve the repair of osteochondral defects in the rabbit knee at early (3 months) and late (12 months) time points.

## METHODS

### Study Design

#### In vivo experiments

Osteochondral defects were surgically created in the knees of skeletally mature New Zealand White Rabbits. Depending on the experiment, the defects were left untreated (empty defects, controls) or filled with clotted bone marrow, clotted bone marrow containing Ad.TGF-β_1_ (BMC+ Ad.TGF-β_1_) or adenovirus carrying the enhanced green fluorescent protein (GFP) cDNA (BMC+Ad.GFP), fibrin hydrogel or fibrin hydrogel containing Ad.TFG-β_1_.

Rabbits receiving Ad.GFP were euthanized after two weeks. Synovium, adjacent cartilage, and tissue from within the osteochondral defect was harvested and transgene expression verified by fluorescence microscopy. Groups of rabbits receiving Ad.TFG-β_1_ were euthanized after 3 months and 12 months. The distal femurs were stained with safranin orange-fast green and repair of the osteochondral defects assessed in a blinded fashion by four individuals using a modified O’Driscoll score. Sections at 12 months were further stained for type I and type II collagen by immunohistochemistry.

#### In vitro experiments

Bone marrow was aspirated from the iliac crests of rabbits and aliquots distributed among wells in a 96-well plate. Saline or Ad.TGF-β_1_ were added and mixed with the marrow, which was then allowed to clot. The marrow clots were cultured in well plates for 15 days. Conditioned media were assayed for TGF-β_1_ expression by ELISA. After 28 days the clots were examined by histology for evidence of chondrogenesis.

Fibrin gels were formed in 96-well plates. Saline, Ad.TGF-β_1_ or Ad.GFP were incorporated into the gel as it formed. Gels were then co-cultured with human MSCs in growth medium. Two days later media were changed to either complete chondrogenic medium or incomplete chondrogenic medium. Transgene expression was assessed by immunofluorescent identification of GFP+ cells and by ELISA measurement of TGF- β _1_ in conditioned media.

#### Adenovirus vector preparation

First generation adenovirus vectors, serotype 5, carrying TGF-β_1_ (Ad.TGF-β_1_) or GFP (Ad.GFP) cDNA under the transcriptional control of the CMV promoter were produced in 293 cells as previously described.^19^ Cell lysates were purified by density gradient ultracentrifugation and dialyzed against 10 mM Tris-HCl, pH 7.8, 150 mM NaCl, 10 mM MgCl_2_ and 4% sucrose. Aliquots were stored at −80°. Titer was estimated as 4 × 10^12^ viral particles (vp) ml^−1^ by OD 260.

#### Marrow clots

Bone marrow (2 ml) was aspirated from the iliac crest of an anesthetized rabbit using a 16-gauge needle. This was distributed in 250 μl aliquots among wells in a 96-well plate. After adding 25 μl saline or adenovirus suspension (10^9^ vp) the marrow was titrated and then allowed to clot for 20 min, by which time the clot was firm enough to enable handling (figure 1).

**Figure 1:**
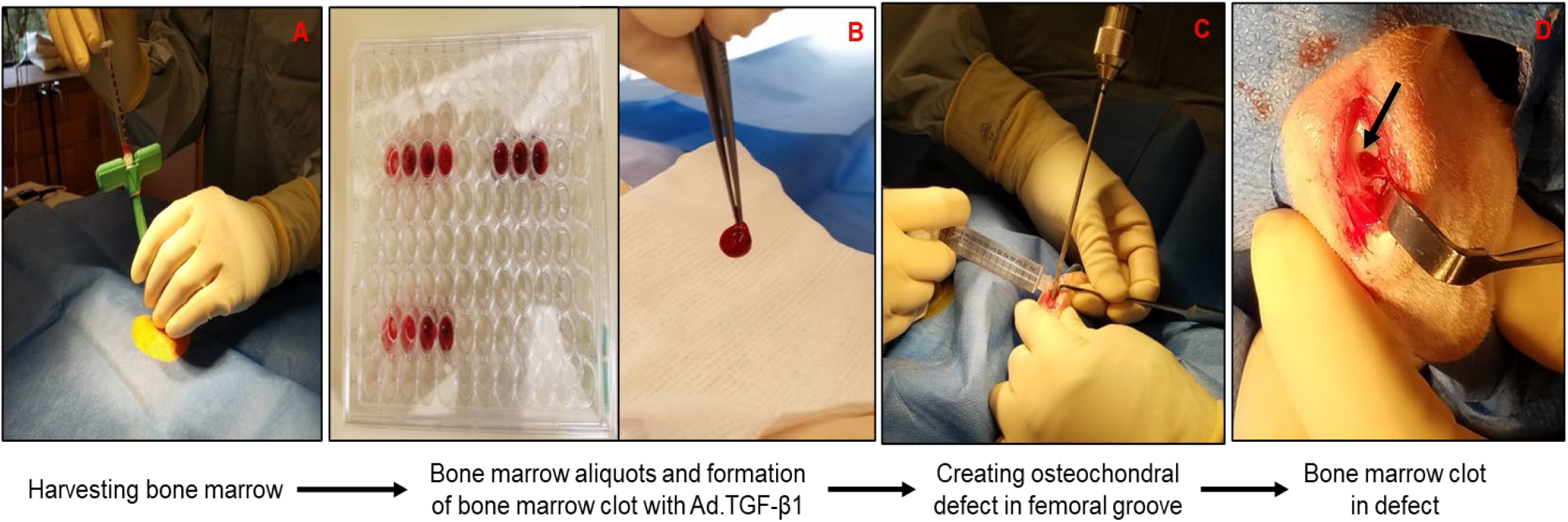
Summary of surgical procedures. Bone marrow was harvested from the iliac crest of the New Zealand white rabbits (A) and 250 μL aliquots of marrow were dispensed into a 96-well plate (B); 25 μL of Ad.TGF-β_1_ were added and the mixture titurated. After approximately 15-20 min, the bone marrow aliquots coagulated and formed into a gelatinous plug, which could be picked up with sterile forceps (B). An osteochondral defect (3.2 mm wide and 5-8 mm deep) was created in the patellar groove (C) and the bone marrow clot was press-fit into the defect (D).

Implantation of bone marrow clots (BMC) into osteochondral defects is described in the section on rabbit surgery. For *in vitro* experiments, BMCs were transferred to Ultra Low-Attachment 24-well plates (Corning, Corning, NY) and cultured in 1 ml incomplete chondrogenic medium (DMEM-HG, 10% FBS, 100 nm dexamethasone, and 1% penicillin/streptomycin) for 28 days with change of medium every 2 or 3 days. The TGF-β_1_ concentration in the conditioned medium was measured by ELISA (R&D Systems, Minneapolis, MN). At day 28 samples were processed for histology using safranin orange and fast green staining.

#### Fibrin hydrogels

Fibrin hydrogels were formed in 96-well plates using Tisseel fibrin glue kit (Baxter Healthcare Corporation, Deerfield, IL). The following ingredients were added in order: 125 μl PBS, 25 μl Ad.GFP or Ad.TGF-β_1_ (10^9^ vp) or PBS, 50 μl thrombin, 150 μl fibrinogen. After 15-20 min at room temperature the polymerized gels were sliced into 4 individual discs and placed in wells on a 96-well plate. To each well were added 250,000 human MSCs (P3-P5, Lonza Ltd., Basel, Switzerland) and incubation continued at 37°C, 5% CO_2_ for 2 days. At this time media were changed to incomplete chondrogenic medium and incubated for a further 16 days, with medium change every 2 days. The TGF-β_1_ concentrations of the media were measured by ELISA and GFP expression observed by fluorescence microscopy. Samples were processed for histology using safranin orange - fast green staining.

#### Osteochondral defect surgery

A total of 15 (n=30 total knees) skeletally mature male New Zealand white rabbits (average weight: 3.54±0.29 kg; average age:9.1±0.6 months) were injected with ketamine (35 mg/kg) and xylazine (5 mg/kg) intra-muscularly and given buprenorphine (0.18 mg/kg) for pre-emptive analgesia. Both hindlimbs and the lumbar region were shaved and the skin sterilized. Rabbits were then intubated with a size 2.5 – 3.5 endotracheal tube and maintained on oxygen and 2-2.5% isoflurane during surgery.

If bone marrow aspirates were required, rabbits were placed in the prone position in a sterile field exposing the lumbar region and a 1 cm incision was made exposing the posterior iliac crest after soft tissue dissection. The iliac crest was penetrated with a bone marrow biopsy needle and the bone was used to aspirate marrow was aspirated from the iliac crest. After the skin closure was closed, rabbits were turned to a supine position in order to create bilateral osteochondral defects within the patellar groove. After disinfection and draping, a 3-cm medial anterior parapatellar incision was made, and the knee joint accessed by opening the joint capsule medial to the patella. The knee was extended and the patella dislocated laterally giving access to the patellar groove. The knee was then flexed and a full-thickness osteochondral defect (3.2 mm diameter × 5-8 mm deep) created within the patellar groove with continuous irrigation. After the defect was created, the surgeon checked to verify that the defect did not perforate the metaphysis or extend into the medullary canal. Where necessary, marrow clots or fibrin hydrogels were press-fit into the defects, the patella relocated, and the joint capsule and skin closed using 3-0 and 4-0 resorbable sutures, respectively. A summary of the surgical procedure is shown in figure 1.

#### Histology

Rabbit condyles were fixed in 10% neutral buffered formalin for 15 days and decalcified in unbuffered 10% EDTA pH 7.4 (Mol-decalcifier, Milestone Medical, Kalamazoo, MI) with constant stirring at 37° using a decalcifying microwave apparatus (KOS Histostation. Milestone Medical). Decalcification was evaluated using a digital x-ray cabinet (MX-20, Faxitron Bioptics, Tucson, AZ). Fixed and decalcified specimens were dehydrated through graded alcohols, embedded in paraffin; 5 μm sections were cut using an automatic microtome (HM 355S, Thermo Scientific Kalamazoo, MI) and mounted onto positively charged slides (Superfrost Plus Microscope Slides, Fisher Scientific, Pittsburgh, PA). Safranin orange – fast green staining was performed according to standard protocols.

Histological scoring of the sections by 4 blinded investigators (CVN, CHE, RDLV, MC) using the modified O’Driscoll scoring scale. An unpaired, student’s t-test was performed to determine statistical significance (p<0.05).

#### Dual Immunohistochemistry

Expression of collagen type I and collagen type II was examined by double staining. Briefly, formalin-fixed tissue sections (5μm-thick) were deparaffinized in xylene and rehydrated in an ethanol/water gradient series, then rinsed in phosphate-buffered saline/0.05% Tween-20 (PBS-T) (Sigma; St. Louis, MO). Endogenous peroxidase activity was blocked with 3% hydrogen peroxide for 10 min, followed by treatment with 0.1% hyaluronidase with 0.1% pronase in tris-buffered saline (all from Sigma) in a water bath for 30 min at 37 °C to unmask collagen antigens. Slide sections were washed 3 times in PBS-T for 5 min each. Endogenous biotin activity was blocked by incubating the slides with the avidin and biotin solutions (SP-2001, Vector Laboratories), then washed 3 times in PBS-T for 5 min each. Nonspecific binding was blocked with animal-free blocker (SP-5035; Vector Laboratories, Burlingame, California) for 30 min at room temperature). Slides were incubated with goat polyclonal anti-type I collagen antibody (dilution 1:250), (Cat No. 1310-01, Southern Biotech, Birmingham, AL) for 1 hour at room temperature (RT). Then, the slides were washed 3 times in PBS-T for 5 min each, followed by incubation with biotinylated rabbit anti-goat IgG antibody (PK-6105; Vector laboratories) for 30 min at RT, washed, and incubated with the avidin-biotin-peroxidase complex solution (PK-6105; Vector laboratories) for a further 30 min at RT.

Collagen type I staining was developed with DAB (3,3’-diaminobenzidine) (SK-4103, Vector Laboratories), a horseradish peroxidase (HRP) substrate that yields a brown product. DAB-slides slides were washed; and subsequently incubated with the second primary antibody, a mouse monoclonal to collagen II (dilution 1:200) (Cat No. 11-116B3, Developmental Studies Hybridoma bank, University of Iowa) in a humidified chamber at 4 °C overnight. The next day, slides were washed 3 times in PBS-T for 5 min each and incubated with the biotinylated horse anti-mouse IgG antibody (AK-5002, Vector laboratories) for 30 min, washed, and then treated with avidin-biotin-alkaline phosphatase complex solution (AK-5002, Vector laboratories) for 30 min at RT. Collagen type II staining was developed with red alkaline phosphatase substrate (SK-5105, Vector Laboratories) that yields a magenta precipitate. Nuclei were counterstained with hematoxylin (H-3404-100, Vector Laboratories), then quickly dehydrated in a graded ethyl alcohol series, cleared with xylene, and mounted with xylene-based mounting medium. To verify specificity of the immunolabeling, isotype controls consisting of goat IgG polyclonal (1:250) and mouse IgG1 kappa (1:200) isotype were included in a set of sections.

## RESULTS

### Gene delivery using bone marrow clots for in vivo osteochondral defect repair

After two-weeks, the BMC+Ad.GFP group (n=3 rabbits; n=6 knees) was euthanized and both knees were harvested from each rabbit (figure 2A). The tissue within the defect was removed using sterile forceps and placed into a well of an ultra low-binding, 24-well plate filled with culture media. Fluorescence microscopy was used to detect transgene expression within the tissue. We found GFP expression within the harvested tissue from all 6 knees, although to variable degrees (figure 2B), confirming gene delivery with transgene expression for at least two weeks after surgery. There was no GFP activity in the cartilage surrounding the defect, and there was negligible expression in the synovium (not shown).

**Figure 2:**
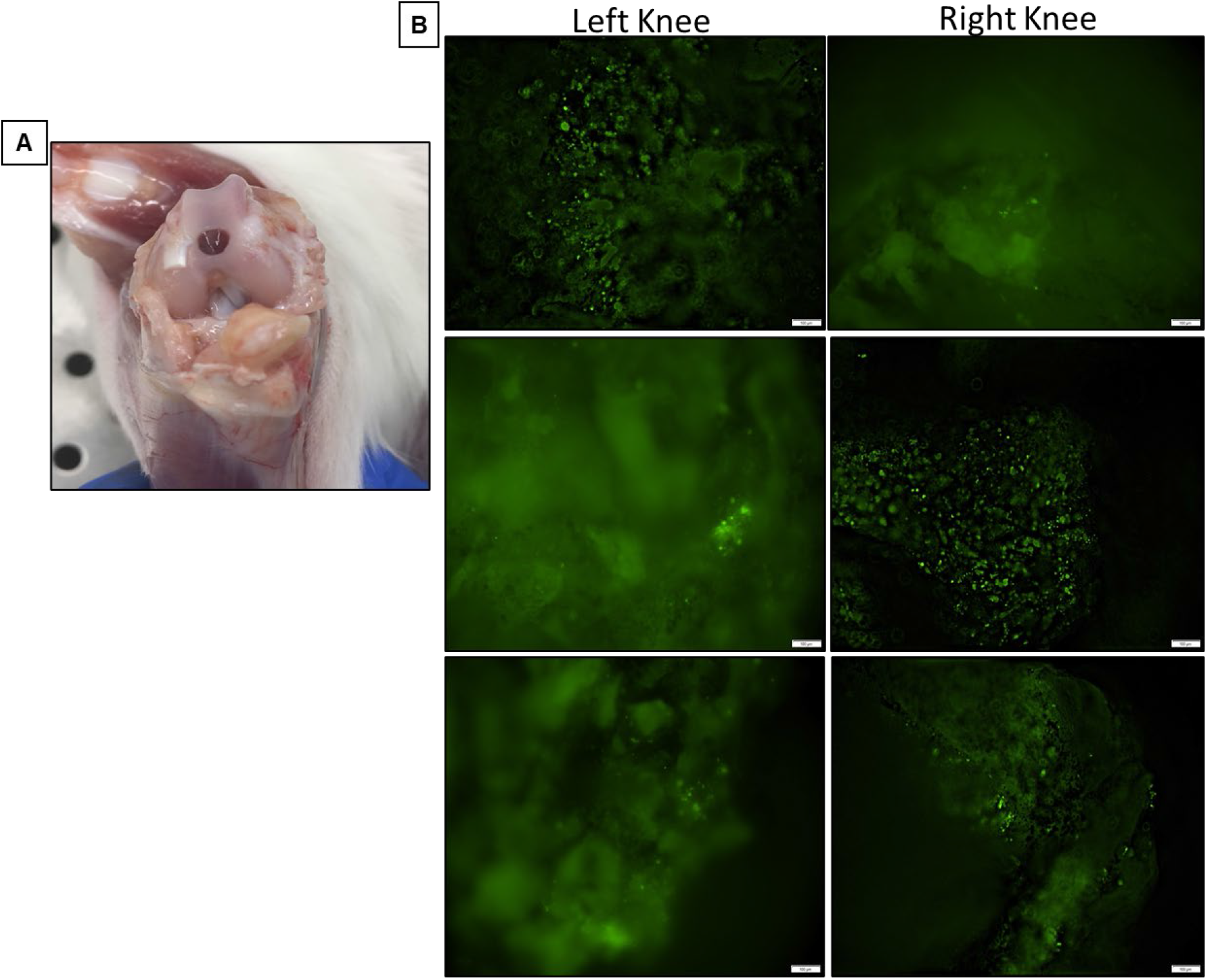
*In vivo* transgene expression. Rabbits in the BMC+Ad.GFP group were sacrificed 2 weeks after surgery and tissue within osteochondral defects (n=3 rabbits; n=6 knees) harvested, GFP expression was confirmed by fluorescence microscopy (A, B).

Clots containing Ad.TGF-β_1_ were implanted into osteochondral defects in the patellar grooves of rabbit knees, with euthanasia at 3 and 12 months after surgery. Both femurs were harvested from each rabbit and processed for histological and immunohistochemistry analysis. Sections through defects in the left and right knees for each rabbit, stained with safranin orange – fast green at 3- and 12-months are shown in figure 3A and figure 4A, respectively. Using a modified O’Driscoll score we found that, despite considerable variability within groups, defects receiving BMC+Ad.TGF-β_1_ had scores that were statistically higher than empty controls at 3 months (figure 3B). These included the overall score, the cartilage component of the score and the osseous component of the score.

**Figure 3:**
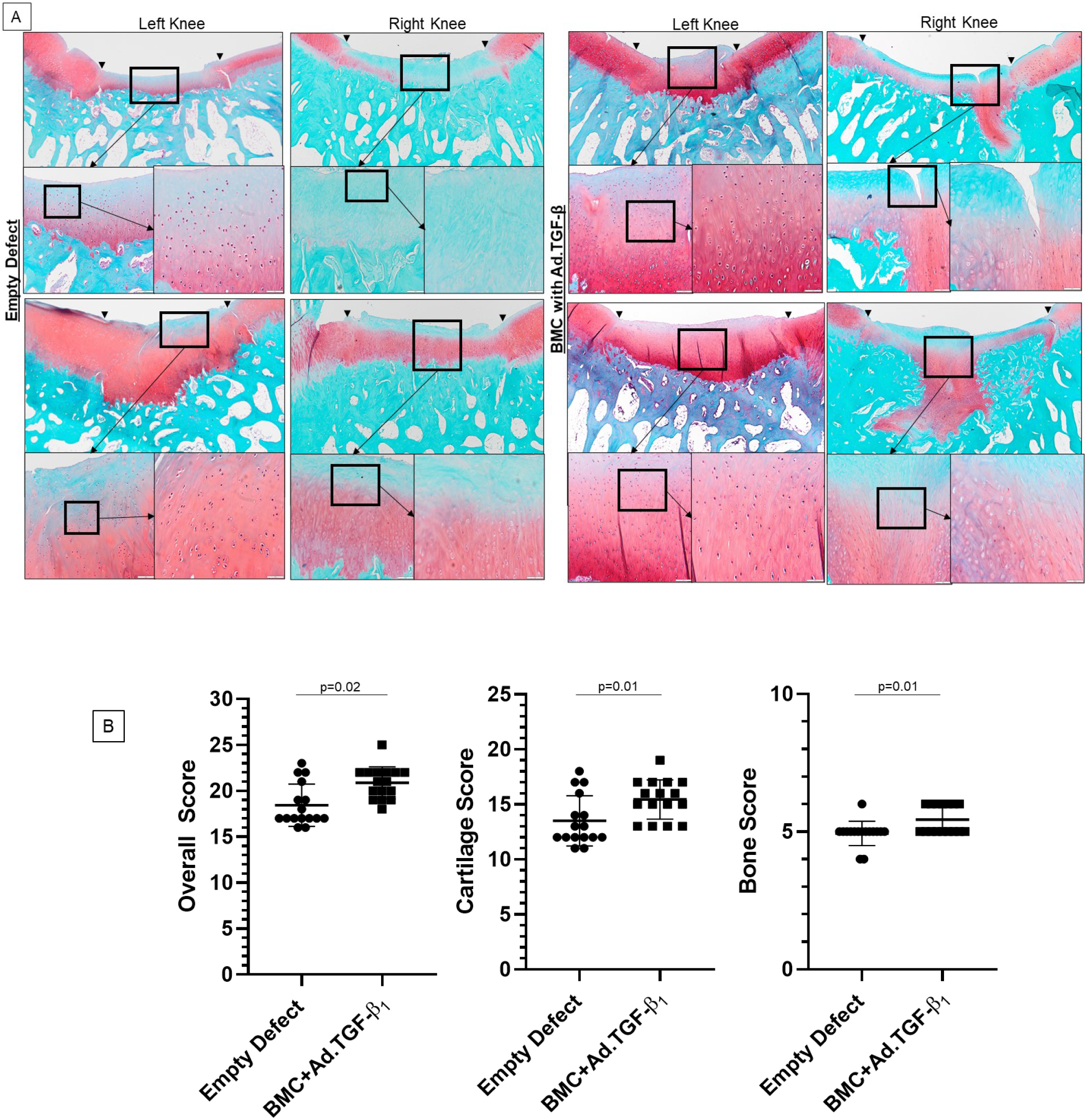
Osteochondral defect repair in empty defect and BMC+Ad.TGF-β_1_ groups at 3 months. Safranin orange - fast green staining of osteochondral defects was used to assess healing in each group (A) (n=4 rabbits). Using the modified O’Driscoll score, knees were scored by four blinded raters who found statistically significant differences in overall scores, cartilage scores and bone scores between the two groups (B).

**Figure 4:**
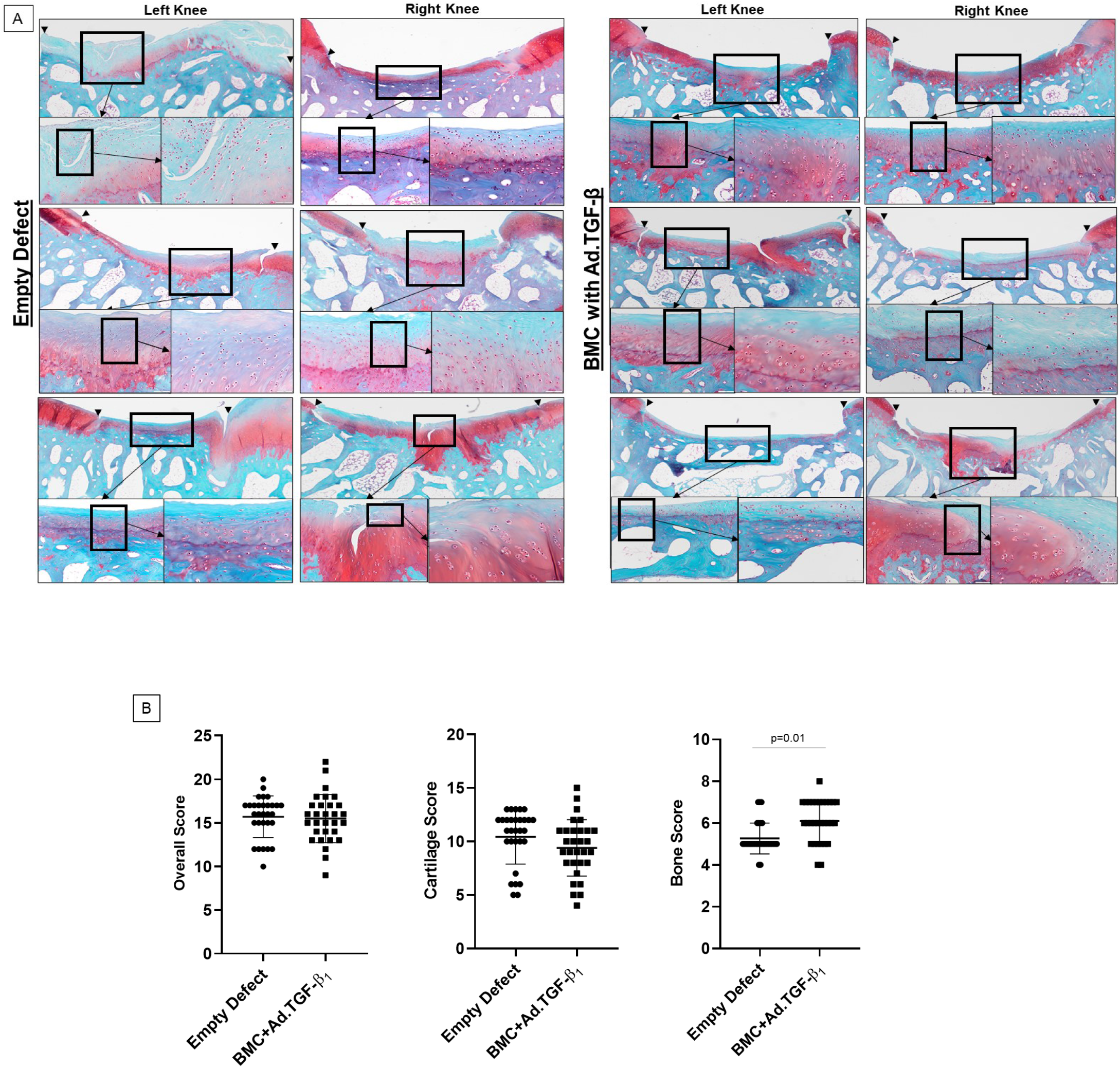
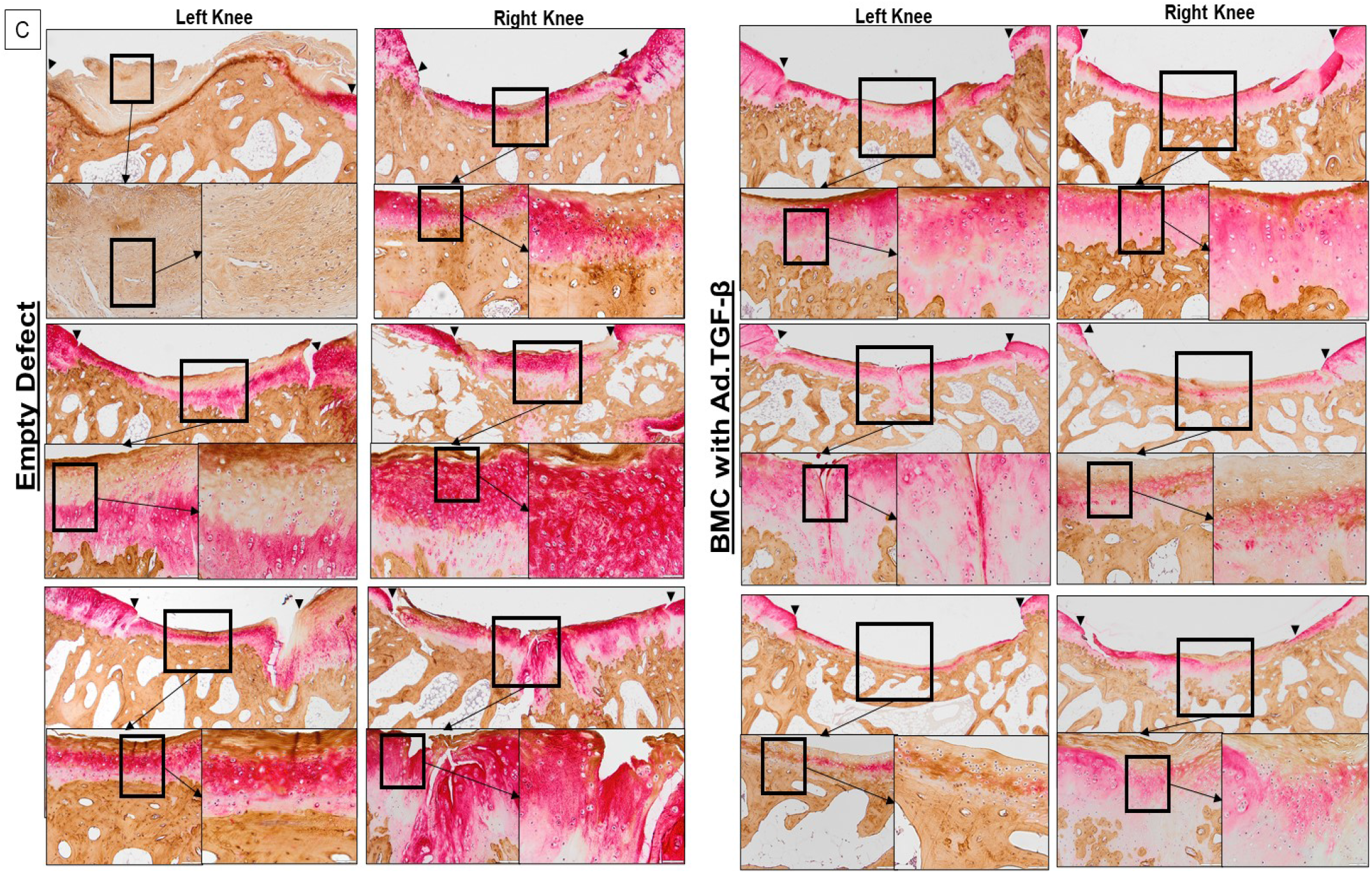
Osteochondral defect repair in the empty defect and BMC+Ad.TGF-β_1_ groups at 12 months. Safranin orange - fast green staining of osteochondral defects was used to assess healing in each group (A) (n=6 rabbits; n=6 knees/group). Using the modified O’Driscoll score, knees were scored by four blinded raters who found no statistically significant differences in overall scores or cartilage scores, but there a significant difference in bone scores between treatment groups (B). Dual immunohistochemistry staining for collagen type I and type II of the osteochondral repair tissue from the empty defect and BMC+Ad.TGF-β_1_ (B) groups one year after surgery was also performed.

However, the differences for overall and cartilage scores disappeared by 12-months after surgery (figure 4B). In all groups there was a decline in overall and cartilage scores by 12-months (p<0.01) such that intergroup differences were no longer apparent. Nevertheless, the bone score continued to improve in the group receiving BMC+Ad.TGF-β_1_ thereby enhancing the statistical significance of the difference (figure 4B).

We also performed dual immunohistochemical staining for collagen type I and type II within the repair tissue of the same groups at 12 months (figure 4C). We found uniform strong staining for collagen-type II within the repair tissue of the BMC+Ad.TGF-β_1_ group, whereas the empty defects stained only weakly for type II collagen.

### Chondrogenic differentiation of bone marrow clots with Ad.TGF-β_1_

Because of the high variability seen in the healing of the Ad.TGF-β_1_ group, we performed a follow-up experiment in which marrow clots were formed with or without Ad.TGF-β_1_ and maintained in culture for 28 days with incomplete chondrogenic medium. During this period, the secretion of TGF-β_1_ was measured by ELISA of conditioned media. After 28 days, clots were prepared for histology with safranin orange-fast green staining. Unmodified, control clots secreted little TGF-β_1_ and did not show any evidence of chondrogenesis (figure 5A, C). Only one of the three clots containing Ad.TGF-β_1_ secreted markedly elevated TGF-β_1_ and had regions of chondrogenesis under histological examination (figure 5B, C).

**Figure 5:**
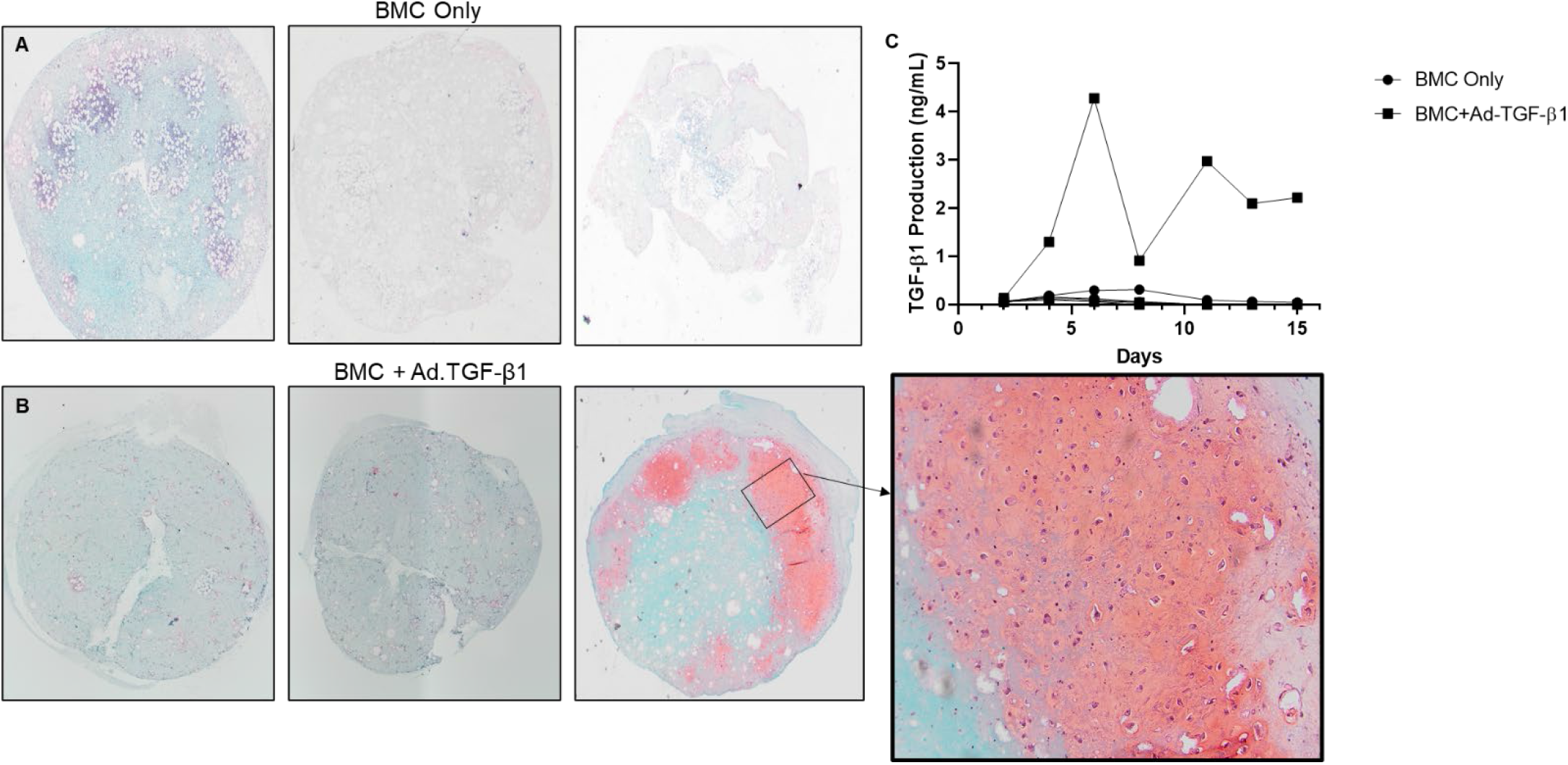
In vitro chondrogenesis of bone marrow coagulates (BMC) with or without Ad.TGF-β_1_ incorporation after 15 days. Safranin-orange staining at day 15 of the BMC only (A) found that none of the clots underwent chondrogenesis while only one of the three clots containing BMC+Ad.TGF (B) had evidence of chondrogenesis showing the presence of chondrocytes surrounded by a rich proteoglycan extracellular matrix. The supernatant from each clot was assayed for TGF-β_1_ (C); the single clot that underwent chondrogenesis was also the only clot producing significant amounts of TGF-β_1_.

### Chondrogenic differentiation of MSCs using fibrin scaffolds incorporating Ad.TGF-β_1_

Much of the variability shown in figures 2-6 may reflect variation in the quality of the bone marrow aspirated from the iliac crests of the rabbits. As a more uniform and reliable alternative to bone marrow, we investigated the use of commercially available fibrin.

**Figure 6:**
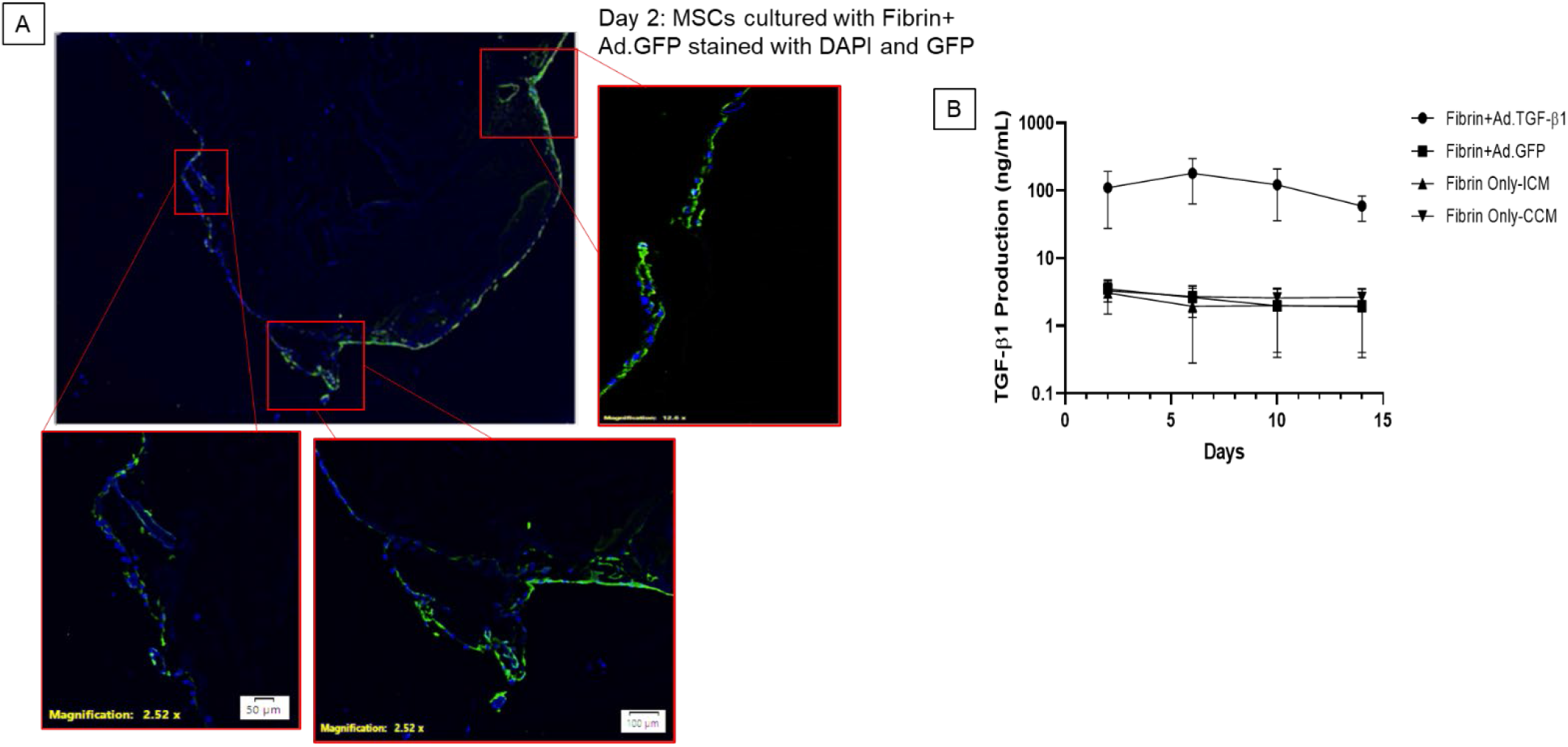
Gene transfer to human MSCs in culture using fibrin scaffolds. MSCs were co-cultured with a fibrin hydrogel incorporating Ad.GFP. By 2 days of co-culture the MSCs were migrating into the scaffold,becoming transduced with Ad.GFP, and expressing GFP (A). We also cultured MSCs with fibrin scaffolds containing Ad.TGF-β_1_ and found that the expression of TGF-β_1_ to be much higher than the control scaffolds lacking Ad.TGF-β_1_ (B).

Human MSCs were co-cultured with a fibrin hydrogel incorporating adenoviral vectors encoding GFP (Ad.GFP). As shown in figure 6A, we observed MSC migration into the scaffold with transduction and expression of GFP after only 2 days of co-culture. When the fibrin hydrogels contained Ad.TGF-β_1_, expression of TGF-β_1_ was very high, exceeding 180 ng/mL at its maximum on day 6. Control groups produced a basal level of approximately 4 ng/mL TGF-β_1_ (figure 6B).

### Gene delivery using fibrin scaffolds for in vivo osteochondral defect repair

With the encouraging *in vitro* data shown in figure 6, a small pilot study explored the use of the fibrin+Ad.TGF-β_1_ scaffold to repair osteochondral defects. Three osteochondral defects received the fibrin+Ad.TGF-β_1_ construct, and one received fibrin alone. After 3-months, the femurs were harvested and processed for histology and immunohistochemistry for collagen type II.

Sections through the middle of the repair tissue were stained with safranin orange-fast green are shown in Figure 7A. The control defect which received the empty fibrin scaffold demonstrated very poor healing. Defects which received the fibrin+Ad.TGF-β_1_ scaffolds had repair tissue that was rich in proteoglycan and resembled articular cartilage; with one exception, was continuous with the surrounding native tissue.

**Figure 7:**
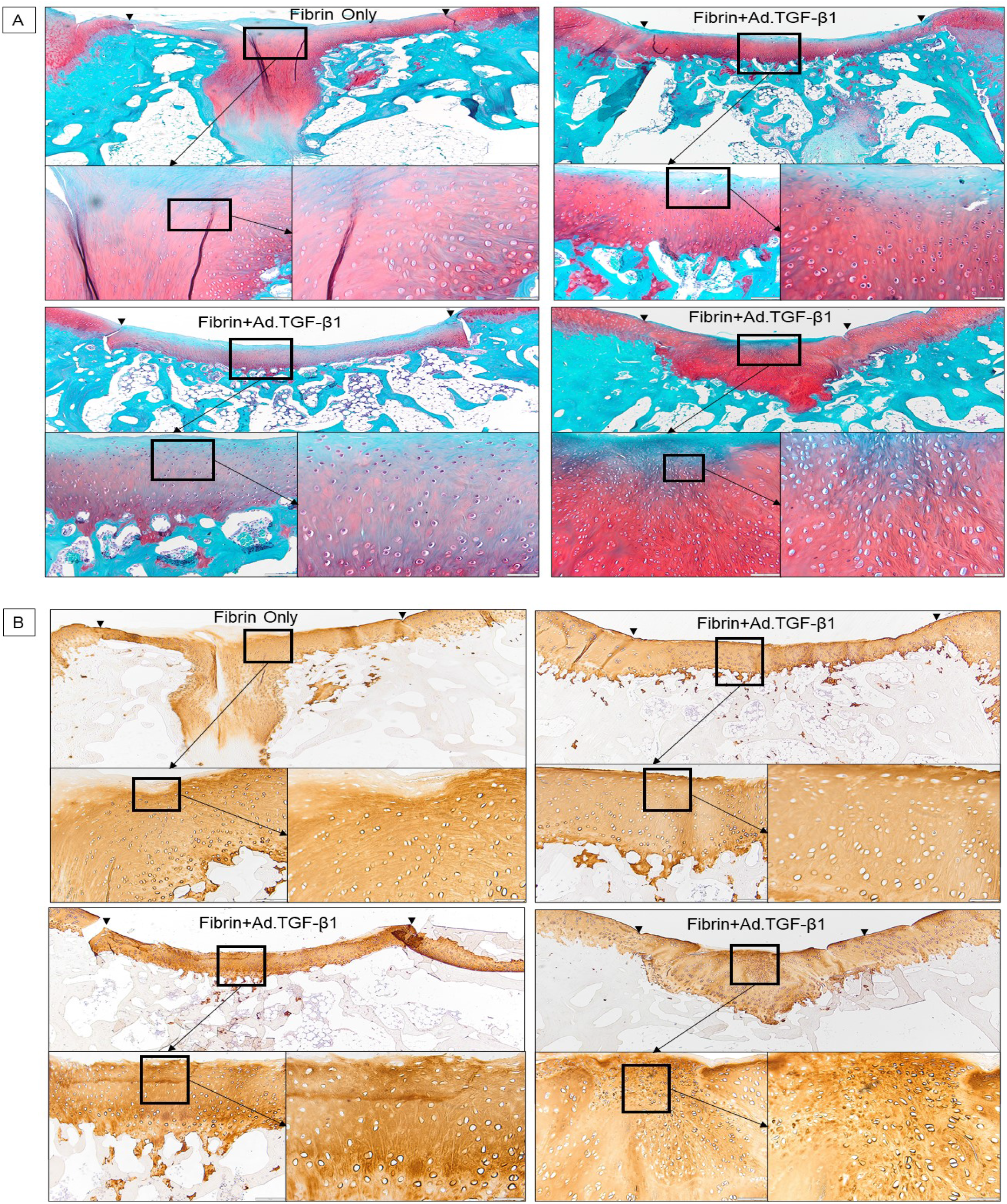
Gene delivery using fibrin scaffolds for in vivo osteochondral defect repair. Safranin orange - fast green staining of histological sections of the osteochondral defects demonstrate differential healing response between defects containing empty scaffolds and those containing Ad.TGF-β_1_ (A). Collagen type II staining of the same sections revealed abundant collagen type II in the repair tissue of the knees which received the fibrin scaffolds with Ad.TGF-β_1_ (B).

We also performed immunohistochemistry for collagen-type II for these knees (Figure 7B). We found strong staining for collagen type II in defects receiving the fibrin+Ad.TGF-β_1_ scaffold whereas defects receiving the control fibrin scaffold had less staining for type II collagen.

## DISCUSSION

Gene transfer offers to enhance the regenerative behavior of musculoskeletal tissues,^20^ including cartilage^21^ while single-step procedures that can be delivered at point-of-care will facilitate clinical translation and utilization.^22^ In the context of cartilage repair, previous research has confirmed that bone marrow clots with embedded adenovirus vectors can deliver transgenes to osteochondral defects, leading to transgene expression within the lesion.^17^ Consistent with this, Sieker et al.^18^ used bone marrow coagulates with adenoviral vectors encoding BMP-2 and IHH in a rabbit osteochondral defect model. Expression of BMP-2 gave encouraging early repair but led to the formation of osteophytes and elevated subchondral bone after 13 weeks.^18^ IHH, in contrast, generated an abundance of immature, highly cellular cartilage that did not remodel into mature articular cartilage during the 13 week experiment.^18^ Ivkovic et al.^23^ also used BMC to deliver TGF-β_1_ cDNA in a sheep chondral defect model. Although areas of the defect showed *de novo* cartilage formation by 6 months complete repair was not achieved, possibly because there was no connection to the underlying bone marrow.

In the present experiments, healing of osteochondral defects in the patellar groove of the rabbit knee proved highly variable, both in empty defects and in defects receiving bone marrow clots incorporating Ad.TGF-β_1_. There are several sources of variability. Because the surgeries were conducted under closely controlled conditions, variability in the healing of empty defects presumably reflects biological variability between individual New Zealand white rabbits which, although inbred, are not syngeneic. Such intrinsic biological variability would have been compounded in defects receiving Ad.TGF-β_1_ by the heterogeneous nature of bone marrow aliquots. This is best illustrated in figure 6, showing that only 1 in 3 marrow clots incorporating Ad.TGF-β_1_ secreted TGF-β_1_ and contained areas of chondrogenesis. This may be attributable to increasing dilution of the marrow by blood as aspiration continues. Nevertheless it is encouraging that, despite this background variability, a statistically significant improvement in healing was evident at 3 months.

It appears that the variability in the quality of different marrow aspirates can be obviated by the use of a fibrin gel where, under in vitro conditions, the production of TGF-β_1_ by human MSCs was high and uniform. Although a fibrin scaffold lacks the chondroprogenitor cells of marrow, the number of such cells in a 250 μl aliquot of marrow is likely to be small and an acceptable compromise in using the fibrin scaffolds to ensure greater uniformity in gene delivery. Moreover, by eliminating the need to harvest marrow, the use of fibrin eliminates one procedure. The small exploratory experiment using fibrin gels in conjunction with Ad.TGF-β_1_ provided encouraging results that merit further development of this technology. The alternative approach of administering vector directly to the marrow as it enters the lesion from the underlying marrow is also showing promising results in animal models.^24, 25^

The overall O’Driscoll score and that of the cartilage component fell between 3 and 12 months. This is reminiscent of the study by Shapiro et al^26^ who noted a similar trend in the spontaneous healing of osteochondral defects in young rabbits. Thus, although transfer of TGF-β_1_ accelerated and improved early healing, it could not overcome the subsequent delayed decline noted in this model. Addressing this problem will be a matter for future research.

## ACKNOWLEDGEMENTS

The authors would like to thank the support of the Department of Comparative Medicine at Mayo Clinic for their support for excellent animal care and communication.

## CONTRIBUTIONS

All authors have made substantial contributions to the study and approved the final submitted manuscript. Nagelli and Evans take full responsibility for the integrity of the research.

Nagelli: Collection and assembly of data, analysis and interpretation of the data, drafting of the article, final approval of the article

De la Vega: Collection and assembly of data, analysis and interpretation of the data, technical support, editing of manuscript

Coenen: Collection and assembly of data, analysis and interpretation of the data, technical support, editing of manuscript

De Padilla: Collection and assembly of data, analysis and interpretation of the data, technical support, editing of manuscript

Panos: Collection and assembly of data, analysis and interpretation of the data, technical support, editing of manuscript

Trovar: Collection and assembly of data, technical support

Müller: Collection and assembly of data, analysis and interpretation of the data, technical support, editing of manuscript

Evans: Conception and design, collection and assembly of data, analysis and interpretation of the data, drafting of the article, final approval of the article

## ROLE OF FUNDING SOURCE

Support for CVN was received from the National Institute of Arthritis and Musculoskeletal and Skin Diseases grant T32AR56950. CHE’s research is supported, in part, by the John and Posy Krehbiel Professorship in Orthopedics. Funding from the Musculoskeletal Regeneration Partnership Fund by Mary Sue and Michael Shannon is gratefully acknowledged.

## CONFLICT OF INTEREST

There are no conflicts of interests for the authors for this study.

## Notes

### Competing Interest Statement

The authors have declared no competing interest.

## REFERENCES

1. Widuchowski W, Widuchowski J, Trzaska T. Articular cartilage defects: study of 25,124 knee arthroscopies. Knee 2007; 14: 177–182.

2. Sophia Fox AJ, Bedi A, Rodeo SA. The basic science of articular cartilage: structure, composition, and function. Sports Health 2009; 1: 461–468.

3. McCormick F, Harris JD, Abrams GD, Frank R, Gupta A, Hussey K, et al. Trends in the surgical treatment of articular cartilage lesions in the United States: an analysis of a large private-payer database over a period of 8 years. Arthroscopy 2014; 30: 222–226.

4. Brittberg M, Lindahl A, Nilsson A, Ohlsson C, Isaksson O, Peterson L. Treatment of deep cartilage defects in the knee with autologous chondrocyte transplantation. New England Journal of Medicine 1994; 331: 889–895.

5. Mistry H, Connock M, Pink J, Shyangdan D, Clar C, Royle P, et al. Autologous chondrocyte implantation in the knee: systematic review and economic evaluation. Health Technol Assess 2017; 21: 1–294.

6. Nho SJ, Pensak MJ, Seigerman DA, Cole BJ. Rehabilitation after autologous chondrocyte implantation in athletes. Clin Sports Med 2010; 29: 267–282, viii.

7. Steadman JR, Rodkey WG, Singleton SB, Briggs KK. Microfracture technique for full-thickness chondral defects: Technique and clinical results. Operative Techniques in Orthopaedics 1997; 7: 300–304.

8. Steadman JR, Briggs KK, Rodrigo JJ, Kocher MS, Gill TJ, Rodkey WG. Outcomes of microfracture for traumatic chondral defects of the knee: Average 11-year follow-up. Arthroscopy - Journal of Arthroscopic and Related Surgery 2003; 19: 477–484.

9. Knutsen G, Drogset JO, Engebretsen L, Grontvedt T, Ludvigsen TC, Loken S, et al. A Randomized Multicenter Trial Comparing Autologous Chondrocyte Implantation with Microfracture: Long-Term Follow-up at 14 to 15 Years. J Bone Joint Surg Am 2016; 98: 1332–1339.

10. Camp CL, Stuart MJ, Krych AJ. Current concepts of articular cartilage restoration techniques in the knee. Sports Health 2014; 6: 265–273.

11. Ogura T, Bryant T, Minas T. Long-term Outcomes of Autologous Chondrocyte Implantation in Adolescent Patients. Am J Sports Med 2017; 45: 1066–1074.

12. Nehrer S, Spector M, Minas T. Histologic analysis of tissue after failed cartilage repair procedures. Clin Orthop Relat Res 1999: 149–162.

13. Mithoefer K, McAdams T, Williams RJ, Kreuz PC, Mandelbaum BR. Clinical efficacy of the microfracture technique for articular cartilage repair in the knee: an evidence-based systematic analysis. Am J Sports Med 2009; 37: 2053–2063.

14. Oussedik S, Tsitskaris K, Parker D. Treatment of articular cartilage lesions of the knee by microfracture or autologous chondrocyte implantation: a systematic review. Arthroscopy 2015; 31: 732–744.

15. Palmer GD, Steinert A, Pascher A, Gouze E, Gouze JN, Betz O, et al. Gene-induced chondrogenesis of primary mesenchymal stem cells in vitro. Molecular Therapy 2005; 12: 219–228.

16. Steinert AF, Palmer GD, Pilapil C, Nöth U, Evans CH, Ghivizzani SC. Enhanced in vitro chondrogenesis of primary mesenchymal stem cells by combined gene transfer. Tissue Engineering - Part A 2009; 15: 1127–1139.

17. Pascher A, Palmer GD, Steinert A, Oligino T, Gouze E, Gouze JN, et al. Gene delivery to cartilage defects using coagulated bone marrow aspirate. Gene Therapy 2004; 11: 133–141.

18. Sieker JT, Kunz M, Weissenberger M, Gilbert F, Frey S, Rudert M, et al. Direct bone morphogenetic protein 2 and Indian hedgehog gene transfer for articular cartilage repair using bone marrow coagulates. Osteoarthritis Cartilage 2015; 23: 433–442.

19. Gouze JN, Stoddart MJ, Gouze E, Palmer GD, Ghivizzani SC, Grodzinsky AJ, et al. In vitro gene transfer to chondrocytes and synovial fibroblasts by adenoviral vectors. Methods Mol Med 2004; 100: 147–164.

20. Evans CH, Huard J. Gene therapy approaches to regenerating the musculoskeletal system. Nat Rev Rheumatol 2015; 11: 234–242.

21. Evans CH, Ghivizzani SC, Smith P, Shuler FD, Mi Z, Robbins PD. Using gene therapy to protect and restore cartilage. Clinical Orthopaedics and Related Research 2000: S214–S219.

22. Evans CH, Palmer GD, Pascher A, Porter R, Kwong FN, Gouze E, et al. Facilitated endogenous repair: Making tissue engineering simple, practical, and economical. Tissue Engineering 2007; 13: 1987–1993.

23. Ivkovic A, Pascher A, Hudetz D, Maticic D, Jelic M, Dickinson S, et al. Articular cartilage repair by genetically modified bone marrow aspirate in sheep. Gene Ther 2010; 17: 779–789.

24. Cucchiarini M, Madry H. Overexpression of human IGF-I via direct rAAV-mediated gene transfer improves the early repair of articular cartilage defects in vivo. Gene Ther 2014; 21: 811–819.

25. Cucchiarini M, Asen AK, Goebel L, Venkatesan JK, Schmitt G, Zurakowski D, et al. Effects of TGF-beta Overexpression via rAAV Gene Transfer on the Early Repair Processes in an Osteochondral Defect Model in Minipigs. Am J Sports Med 2018; 46: 1987–1996.

26. Shapiro F, Koide S, Glimcher MJ. Cell origin and differentiation in the repair of full-thickness defects of articular cartilage. J Bone Joint Surg Am 1993; 75: 532–553.

